# A comprehensive map of preferentially located motifs reveals distinct proximal *cis*-regulatory elements in plants

**DOI:** 10.1101/2022.01.17.476590

**Authors:** Julien Rozière, Cécile Guichard, Véronique Brunaud, Marie-Laure Martin, Sylvie Coursol

## Abstract

The identification of *cis*-regulatory elements controlling gene expression is an arduous challenge that is being actively explored to discover the key genetic factors responsible for traits of agronomic interest. Here, we have used a *de novo* and genome-wide approach for preferentially located motif (PLM) detection to investigate the proximal *cis*-regulatory landscape of *Arabidopsis thaliana* and Zea *mays*. We report three groups of PLMs in each gene-proximal region and emphasize conserved PLMs in both species, particularly in the 3’-gene-proximal region. Comparison with resources of transcription factor and microRNA binding sites indicates that 79% of the identified PLMs are unassigned, although some are supported by MNase-defined cistrome occupancy analysis. Enrichment analyses further reveal that unassigned PLMs provide functional predictions distinct from those inferred by transcription factor and microRNA binding sites. Our study provides a comprehensive map of PLMs and points at their potential utility for future characterization of orphan genes in plants.

## Introduction

Plants, as sessile organisms, need to adapt to local constraints such as bacteria, fungi and pest presence, as well as environmental changes. One of the fundamental drivers of their adaptation is the activation or repression of gene transcription (1–5). These processes are tuned by many regulatory factors, including transcription factors (TFs) that bind to specific DNA TF binding sites (TFBSs) (6–9).

Efforts to characterize TFBSs at the genome-wide scale rely mostly on the *in vivo* chromatin immunoprecipitation followed by sequencing (ChIP-seq) method (10) or the in *vitro* DNA affinity purification (DAP-seq) method (11). Nevertheless, these two TF-centered methods are limited by the resolution of the data (fragments of about 100 bp), the difficulty of analyzing many TFs with specific antibodies (ChIP-seq) or the lack of consideration of potential co-factors and the genomic context (DAP-seq). In the 2020 ReMap update (12), only 372 transcriptional regulators were thus collected out of the 1,533 encoded by *Arabidopsis thaliana* (13). More globally the dependence on experimental conditions does not allow the identification of a complete cistrome for each TF studied. Chromatin structure profiling methods, including assay for transposase-accessible chromatin with high-throughput sequencing (ATAC-seq) (14), DNase I hypersensitive sites sequencing (DNase-seq) (15) or more recently MNase-defined cistrome-occupancy analysis (MOA-seq) (16) also depend on experimental conditions and may present resolution limitations to identify TFBSs.

Alternative and complementary approaches to identify potential TFBSs include *in silico* predictive methods (17–19). These DNA-centered methods have several advantages: they are capable to rapidly analyze a large number of potential TFBSs, identify *de novo* potential *cis*-regulatory elements without *a priori,* produce the highest-resolution footprints of DNA binding sites, and last but not least, are not dependent on experimental conditions. In this regard, PLMdetect (Preferentially Located Motif detection) (20) currently detects known *cis*-regulatory elements that are over-represented at a specific location relative to the transcription start site (TSS) in *A. thaliana* and are therefore referred to as preferentially located motifs (PLMs) (21–24). No large-scale functional PLMdetect-based study has been performed so far on plant proximal regions, including the 3’-untranslated region that also contributes to transcriptional regulation (1, 2, 25). Accordingly, our current understanding towards the global regulatory impact of PLMs remains severely limited.

To this aim, we carried out a genome-wide and *de novo* PLMdetect-based study to comprehensively analyze the 5’- and 3’-proximal regions of genes from *A. thaliana* and *Zea mays.* We sought to determine how their differences in genome content and architecture would be reflected in the characteristics of their PLMs in both proximal regions, including their promoters and untranslated transcribed regions. Our findings revealed the organizing principle of the plant PLM landscape and provide a valuable resource for characterizing unannoted genes.

## Results

### Large-scale PLM identification in gene-proximal regions of *A. thaliana* and *Z. mays*

To define the PLM profile associated with the 5’- and 3-gene-proximal regions, we performed a large-scale and *de novo* PLM detection (Fig. 1). Distribution according to the score values revealed two populations of PLMs with a score below or above 2 in each gene-proximal region of each species (Supplementary Fig. S1). The higher PLM score means that a PLM is highly predictive of a *cis*-regulatory element in a specific context. Subsequently, only the PLM population with a score above 2 was considered, leading to the identification of 6,998 and 9,768 (7,447 and 6,639) PLMs in the 5’ (3’)-gene-proximal regions (referred to as 5’ (3’)-PLMs) of *A. thaliana* and Z. *mays,* respectively (Fig. 2a and Supplementary Table 1). To verify that the PLMs detected were not redundant, we tested the inclusion relationship between two PLMs (a k-mer included in a larger k-mer) when they shared 50% of their functional windows and were present in the same gene sets (Supplementary Table 2). Only two pairs of PLMs corresponding of four 5’-PLMs of Z. *mays* shared the same PLM-containing gene sets. We therefore maintained our initial number of PLMs based on the score for all subsequent analyses.

**Fig. 1.**
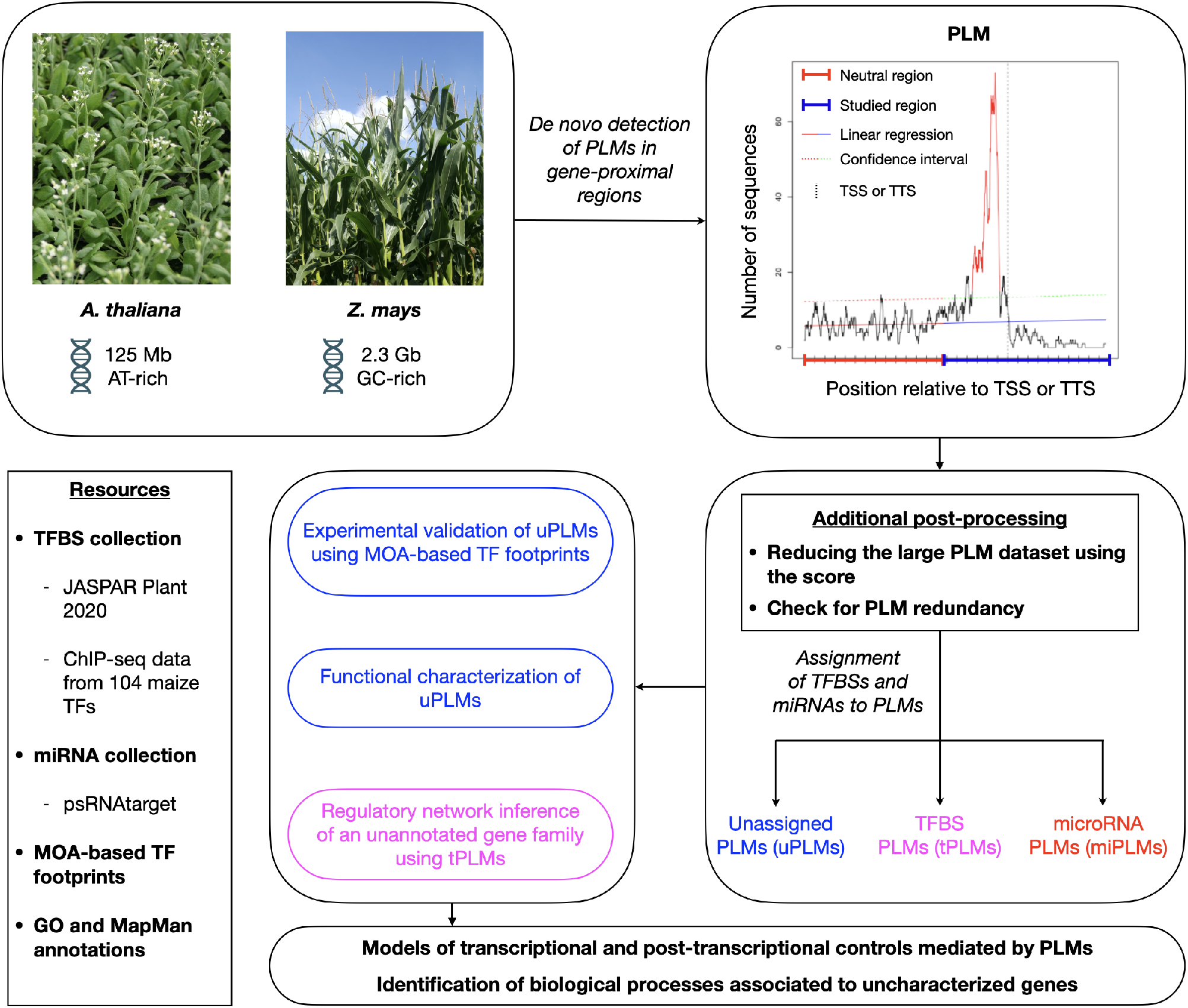
A schematic overviewing the workflow used in this study in *de novo* and genome-wide quantification, characterization and exploitation of PLMs in gene-proximal regions of *A. thaliana* and *Z. mays.* We first detected PLMs in the 5’- and 3’-proximal regions of genes from both plant species. Next, we selected the PLMs according to their score, checked for PLM redundancy and determined whether some of the detected PLMs might be TFBSs (referred to as tPLMs) or targeted by miRNAs (referred to as miPLMs) using distinct resources of TFBSs (JASPAR Plant 2020 (31) and top 1% k-mers from ChIP-seq data of 104 maize TFs (32)) and miRNA binding sites (psRNATarget (37)). Unassigned PLMs (referred to as uPLMs)-containing gene sets were functionally characterized with GO and MapMan enrichments relative to the genome. Moreover, we showed that some 5’-uPLMs were supported by MOA-based TF footprints data (16) of *Z. mays.* Using 5’-tPLMs, we finally inferred the regulatory network of a poorly characterized maize-specific gene family. Collectively, our results provide functional inside into the regulatory role of PLMs and highlight biological processes associated to uncharacterized genes.

**Fig. 2.**
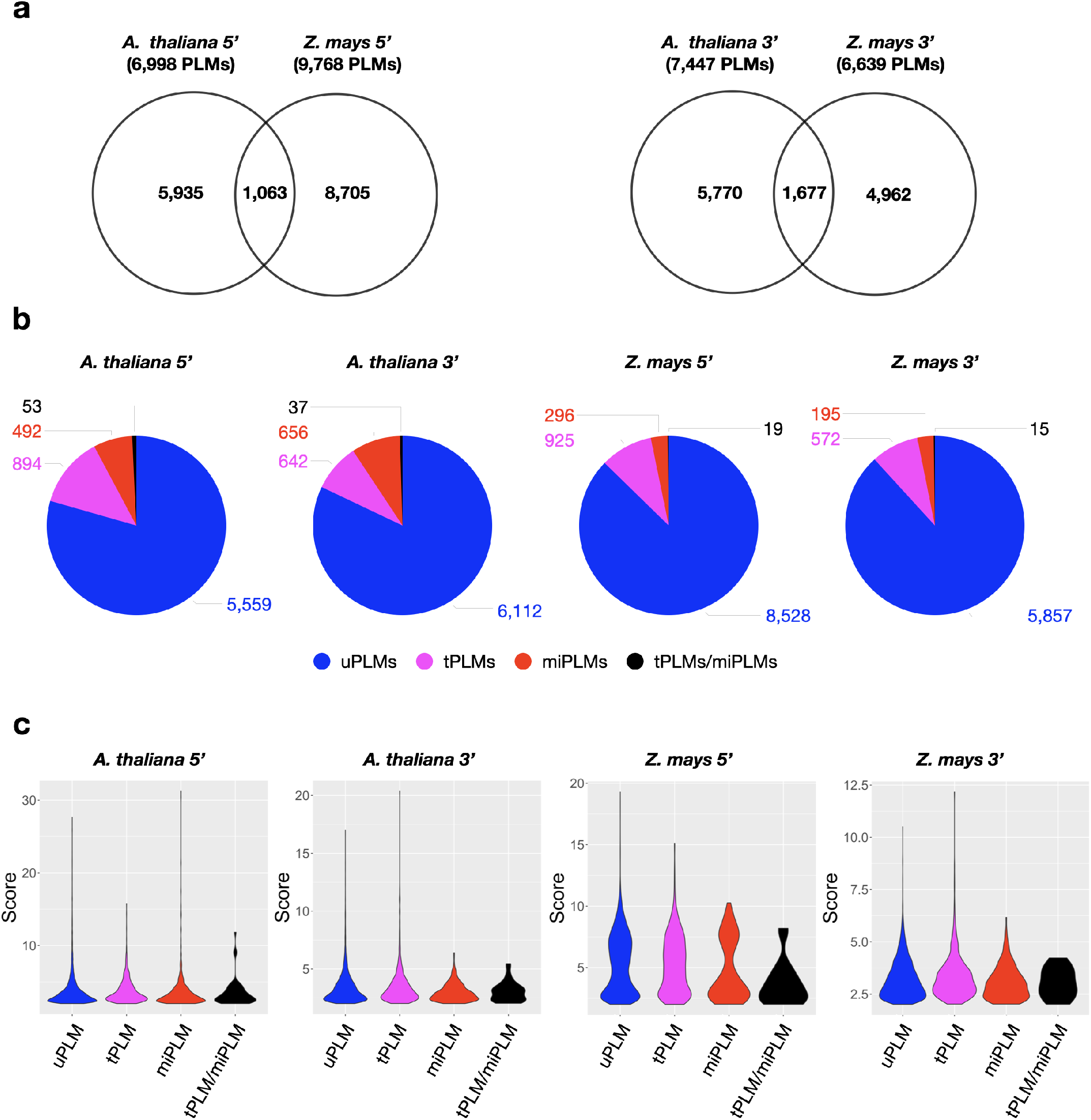
Characterization of PLM content in gene-proximal regions of *A. thaliana* and *Z. mays*. **(a)** Venn diagram showing the extent of overlap between 5’- or 3’-PLMs of *A. thaliana* and *Z. mays.* **(b)** Dissection of PLM types identified in the 5’- or 3’-gene-proximal region of *A. thaliana* and *Z. mays.* **(c)** Violin plot of PLM scores according to the PLM types in the 5’- or 3’-gene-proximal region of *A. thaliana* and *Z. mays.*

Comparison of the PLM content of the two species revealed that *A. thaliana* and Z. *mays* shared 1,063 5’-PLMs and 1,677 3’-PLMs (Fig. 2a). It is worth noting that 98% of these PLMs were located in the 200 bases surrounding the TSS or the transcription termination site (TTS). Inspection of the preferential position of the PLMs further revealed three distinct groups within each targeted region of each species with similar distribution patterns (Fig. 3). In the 5’-gene-proximal region, groups 1 and 2 were localized upstream of the TSS, while group 3 was localized on the TSS. In the 3’-gene-proximal region, groups 1 and 2 were localized upstream of the TTS, while group 3 was localized on the TTS. We did also observe that 5’-PLMs in *Z. mays* were more spread out along the sequence than those of *A. thaliana.*

**Fig. 3.**
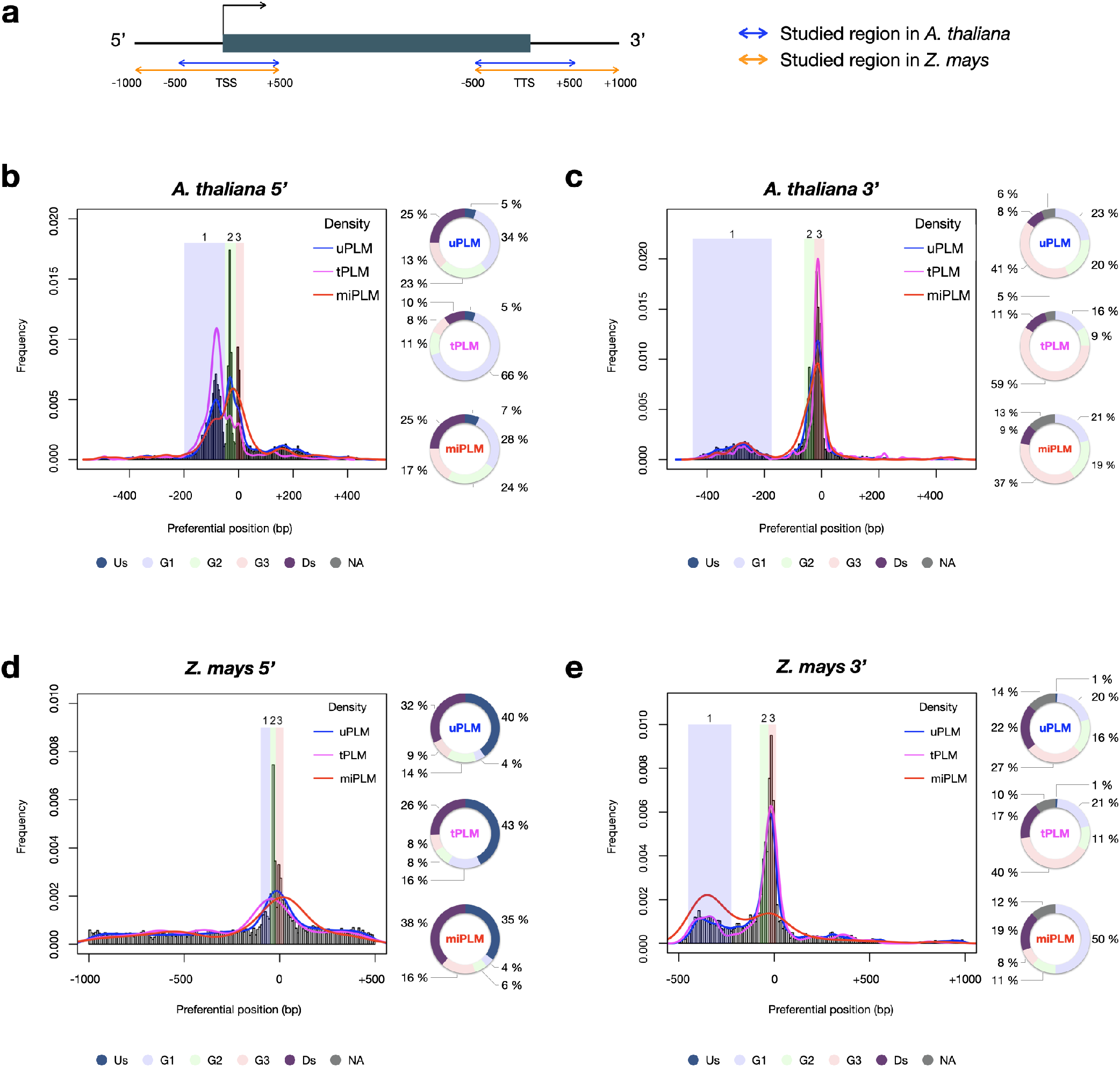
PLM frequency and characterization according to their preferential position. (**a**) Schema of the studied regions with respect to the gene in *A. thaliana* and *Z. mays.* (**b**) 5’-PLMs in *A. thaliana.* Group 1: [-200;-50[; Group 2: [-50;-10[; Group 3: [-10;+20[. (**c**) 3’-PLMs in *A. thaliana.* Group 1: ]-450;-175]; Group 2: ]-60;-25]; Group 3: ]-25;+10]. (**d**) 5’-PLMs in *Z. mays.* Group 1: [-100;-50[; Group 2: [-50;-20[; Group 3: [-20;+20[. (**e**) 3’-PLMs in *Z. mays.* Group 1: ]-450;-225]; Group 2: ]-75;-30]; Group 3: ]-30;+10 bp]. Us: region located upstream the three groups; G1, G2 and G3: groups 1,2 and 3; Ds: region located downstream the three groups; NA: region located between the groups when they are not juxtaposed.

Additionally, each group had specific nucleotide content. Group 1 was composed of A, T, C and G nucleotides in equal proportions in both species. In contrast, group 2 was composed predominantly of A/T (74% and 64% in *A. thaliana* and *Z. mays*, respectively) in agreement with previous observations reporting TATA and TATA-like boxes in this region (26). In the case of group 3, we found that the GC content of the 5’-PLMs differed between the two species (37 % of GC in *A. thaliana vs* 55% in *Z. mays*), in agreement with the report of GC-rich genes in monocot species (27, 28) and recent promoter comparisons using *A. thaliana* and the three cereal species brachypodium, wheat and barley (29).

We did also observe that 3,877 and 2,130 3’-PLMs detected in the [-40;+10] bp interval relative to the TTS (which corresponds to the end of group 2 and the whole group 3) in *A. thaliana* and *Z. mays*, respectively, showed similarities to the *cis*-elements that guide the cleavage and polyadenylation molecular complex (CPMC) essential for mRNA biogenesis (30). These 3’-PLMs were A/T-rich (over 78%) consistently with the far upstream element (FUE) and near upstream element (NUE). Furthermore, those localized 10 bases upstream and downstream of the TTS were composed of sequences rich in T (42% in both species) *>* A (38% in *A. thaliana* and 34% in *Z. mays) >* C (11% in *A. thaliana* and 14% in *Z. mays*) *>* G (9% in *A. thaliana* and 11% in *Z. mays*), in agreement with the known proportions of nucleotides in the cleavage element (CE) (30).

### Identification and positional distribution of TFBS-like PLMs in gene-proximal regions

We anticipated that some of the detected PLMs might be TFBSs. To test this, we used the JASPAR Plant 2020 database (31) combined with the top 100 k-mers from ChIP-seq data of 104 maize TFs (32) and found that 13.5% of 5’-PLMs (9.1% of 3’-PLMs) of *A. thaliana* indeed showed similarities to TFBSs and were therefore referred to as tPLMs (Fig. 1, Fig. 2b and Supplementary Table 3a-3b). In *Z. mays,* 9.6% of 5’-PLMs and 8.8% of 3’-PLMs corresponded to tPLMs (Fig. 2b and Supplementary Table 3c-3d). It is worth noting that, in *A. thaliana*, 5’-tPLMs were more localized in group 1, whereas 3’-tPLMs were more localized in group 3 (Fig. 3b and 3c). In *Z. mays*, 5’-tPLMs were more localized upstream and downstream of the identified groups, while 3’-tPLMs followed the same behavior as in *A. thaliana* with greater localization in group 3. Overall, these results reveal a strong localization of 5’-tPLMs in the interval between 200 and 50 bp upstream of the TSS in agreement with previous observations in *A. thaliana* and *P. Persica* (33, 34). In contrast, 3’-tPLMs mainly localized in the TTS region in both species.

We also investigated how the different TF families were distributed in each proximal region. Among the 47 TF families listed in our reference, 39 and 40 (35 and 37) were susceptible to bind to 5’ (3’)-tPLMs in *A. thaliana* and *Z. mays,* respectively (Fig. 4). We observed that all TF families associated to 3’-PLMs also targeted 5’-PLMs. Furthermore, the SWIM-type zinc fingers, CAMTA, LHY and ARR were specific to the 5’-tPLMs in *A. thaliana,* while the HSF factors, E2F, and BBR-BPC were specific to the 5’-tPLMs in *Z. mays.* Comparing the two species also revealed one (CAMTA) and two (HSF factors and E2F) TF families specifically associated with 5’-tPLMs in *A. thaliana* and *Z. mays,* respectively. For the 3’-gene-proximal region, the BBR-BPC TFs were specific to *A. thaliana*, while the SWIM-type zinc fingers, LHY and ARR TFs were specific to *Z. mays*. Additionally, all TF families were not similarly distributed in each gene-proximal region. For example, the MYB TFs had tPLMs in all three groups of each region and species studied (Fig. 4). In contrast, the Trihelix TFs was only present in group 1 in the 5’-proximal region of *A. thaliana.* Other TF families, such as the G2-like TFs, were likely to target different number of PLM groups according to the region and species considered.

**Fig. 4.**
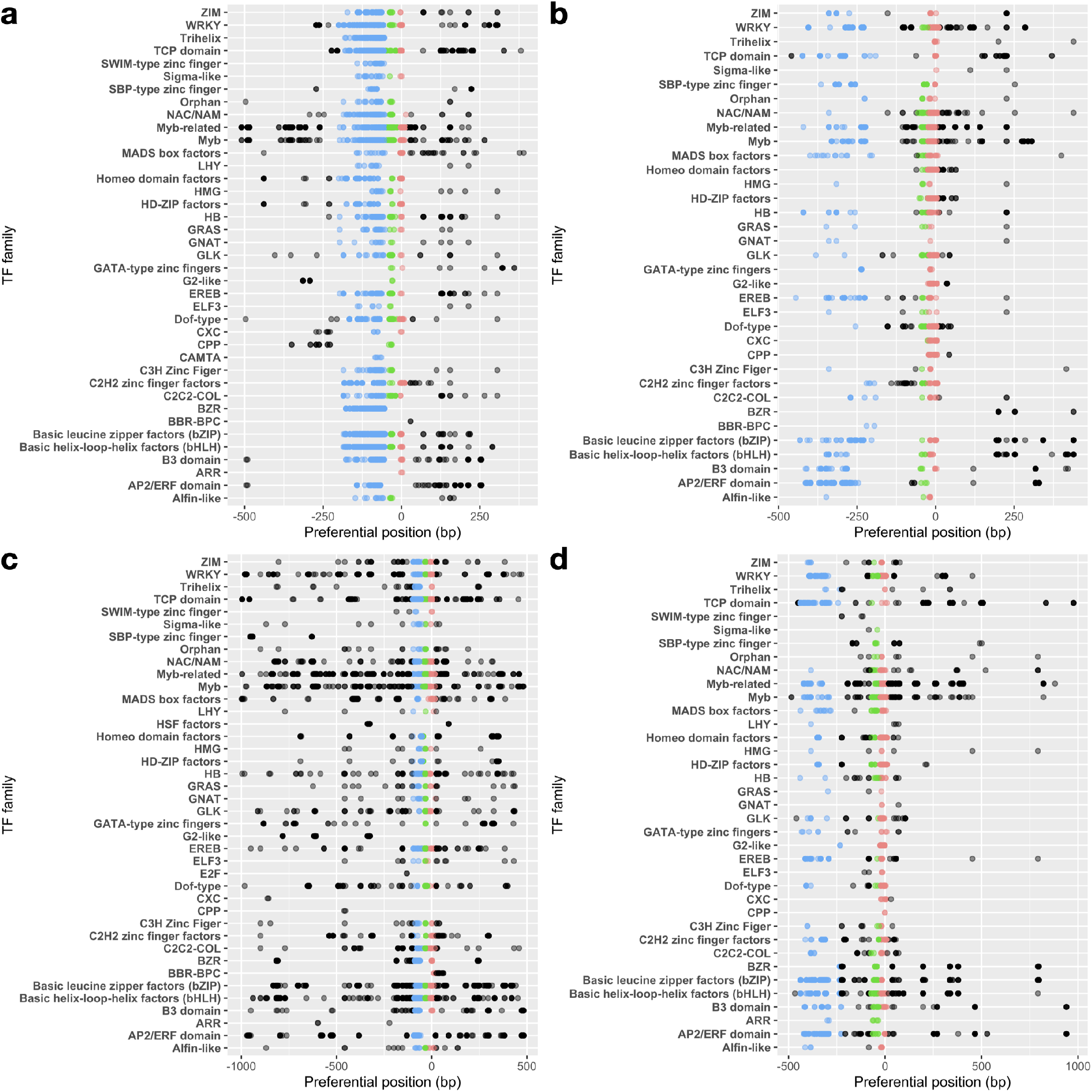
tPLM preferential positions per TF family. tPLMs in 5’- (**a**) and 3’- (**b**) gene-proximal regions of *A. thaliana.* tPLMs in 5’- (**c**) and 3’- (**d**) gene proximal regions of *Z. mays.* Blue, green and red points correspond to the tPLMs in groups 1,2 and 3, respectively. The black points correspond the tPLMs that do not belong to any group. The opacity of the points is relative to the number of tPLMs at that position.

### PLMs occur at miRNA binding sites in gene-proximal regions

Previous studies showed that miRNAs can target transcripts with sequence complementarity (35), thus inducing their degradation. It was also described in *Brassica* that miRNA methylates the promoter region of *SP11* gene to silence it (36). Hence, we predicted that 5’- and 3’-PLMs could be associated to miRNA binding sites. Using the plant small RNA target analysis server psRNATarget (37), we found that 7.8% and 3.2% (9.3% and 3.2%) of the 5’ (3’)-PLMs can be targeted by miRNAs (referred to as 5’ (3’)-miPLMs) in *A. thaliana* and Z. *mays,* respectively (Fig. 1, Fig. 2b and Supplementary Table 4). We noticed that 5’-miPLMs had a maximum density downstream of the TSS, which is consistent with the main mode of action of miRNAs and supports our approach and findings (Fig. 3b and 3d). Surprisingly, more than half of the 5’-miPLMs of *A. thaliana* were located in groups 1 and 2 (Fig. 3b), while those of *Z. mays* were overwhelmingly found outside the groups (Fig. 3d). We also noticed that 3’-miPLMs were more localized in group 3 than in the other two groups in *A. thaliana* (Fig. 3c), while half of them were found in group 1 in *Z. mays* (Fig. 3e).

Next, we investigated which sequence of miPLMs was homologous to that of miRNAs, knowing that the latter are composed of different parts that do not all have the same role (38). We found that 5’-miPLMs of *A. thaliana* showed more frequent homologies to the 5’ seed region (positions 2-8), whereas those of *Z. mays* showed more frequent homologies to the compensatory 3’ end (Fig. 5). For 3’-miPLMs, homologies were more frequent in the center of the miRNA, surrounding the cleavage site in both species, suggesting that miPLMs have distinct functions depending on the region and species considered.

**Fig. 5.**
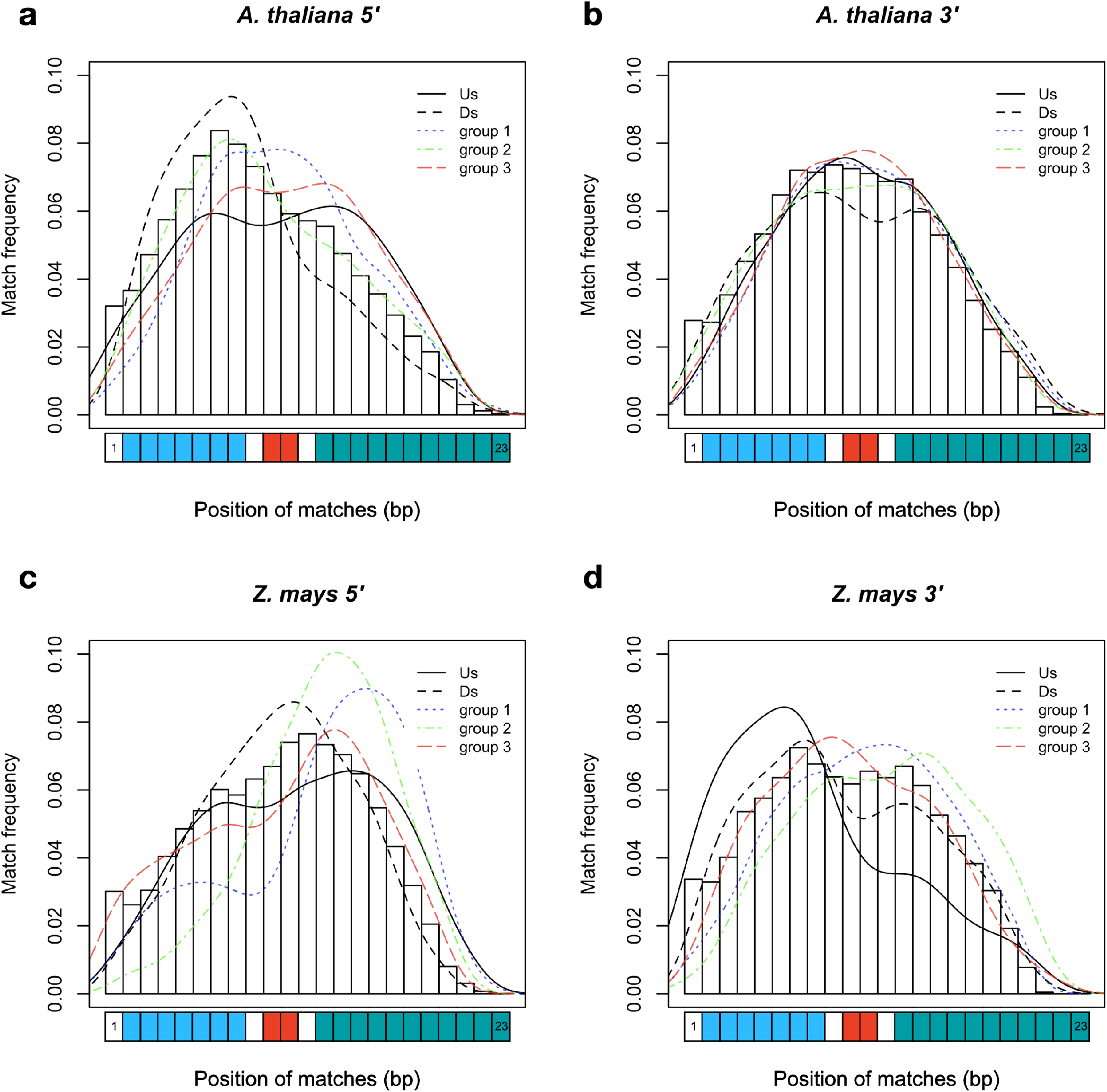
Frequency of miRNA bases covered by miPLMs in gene-proximal regions of A. *thaliana* and Z. *mays*. The color curves indicate the densities of matched-ribonucleotide positions depending on whether the miPLM that matches belongs to one of the PLM groups (groups 1,2 or 3) or is located upstream (Us) or downstream (Ds) of these groups. The blue, red and green rectangles on the abscissa represent the bases of the 5’-seed region, the cleavage site and the 3’-compensatory end of the miRNA, respectively.

Interestingly, we found that 53 and 19 (37 and 15) 5 (3’)- miPLMs corresponded to tPLMs in *A. thaliana* and *Z. may*s, respectively (Fig. 2b and Supplementary Table 1a-1d). In *A. thaliana*, we noticed that WRKY, Basic helix-loop-helix factors (bHLH) and Basic leucine zipper factors (bZIP) represented the three major TF families identified in the 5’-gene-proximal region, while C2H2 zinc finger factors, Myb-related and HD-ZIP factors were the three major TF families identified in the 3’-gene-proximal region (Supplementary Table 4e and 4f). In *Z. mays*, bHLH, bZIP and BZR represented the three major TF families identified in the 5’-gene-proximal region, while bHLH, bZIP and TCP domain were the major TF families in the 3’gene-proximal region (Supplementary Table 4g and 4h).

### Unassigned PLMs include *cis*-regulatory players

Comparison with resources of TF and miRNA binding sites revealed that more than 79% of the identified PLMs were unassigned PLMs (referred to as uPLMs) (Fig. 1, 2b–2c and Supplementary Table 1). This observation prompted us to evaluate the uPLMs’ score and the RNA polymerase II binding site content. We found that 39.2% and 83.1% of the most relevant 5’-uPLMs (score greater than 10) detected in *A. thaliana* and *Z. mays*, respectively, were distinct to RNA polymerase II binding sites (11.5% and 6.9% of the 5’-uPLMs detected in *A. thaliana* and *Z. mays*, respectively) (31), suggesting that uPLMs could include *cis-* regulatory players (Fig. 2c and Supplementary Table 5).

In *A. thaliana,* one-third of the 5’-uPLMs were localized in group 1, while more than one-third of the 3’-uPLMs were localized in group 3. In *Z. mays,* the 5’-uPLMs were preferentially localized upstream and downstream of the three groups detected, showing a greater dispersion than that observed in *A. thaliana.* Furthermore, the 3’-uPLMs of *Z. mays* had a more balanced distribution among the different groups and downstream part than those of *A. thaliana*. Interestingly, the density of the 5’-uPLMs in both species was higher in the core promoter region corresponding to group 2 and known to be the locus of many regulatory events (21, 39–41), confirming the relevance of our hypothesis (Fig. 3).

Recently, a MOA-seq strategy on developping maize ears led to the identification of 215 small (< 30 bp) TF footprints collectively distributed over 100,000 non-overlapping binding sites accross the genome (16). Given the relatively small size of these footprints and their remarkable clustering within the 100 bp proximal to the promoters, we examined them in terms of sequence and position (Fig. 1). We found that 85 of these 215 TF footprints significantly matched with 203 of our motifs. Considering the position of these motifs (plus or minus 50 bases upstream and downstream of the corresponding PLM functional window), 30% of them covered 94 PLMs (Supplementary Table 9), including 24 tPLMs, 14 miPLMs and 60 uPLMs. Overall, these results supports our hypothesis that 5’-uPLMs include *cis*-regulatory players.

### uPLMs provide specific functional predictions

To characterize further the uPLMs, we used GO-term and MapMan functional category enrichment analysis to classify them according to the genes in which they occur (Fig. 1). In both species, the 5’- and 3’-uPLMs-containing gene sets constituted two highly differentiated populations in terms of their biological processes or MapMan categories relative to the other identified PLM classes, further confirming that they include *cis*-regulatory players (Fig. 6 and Supplementary Table 6 and 7).

**Fig. 6.**
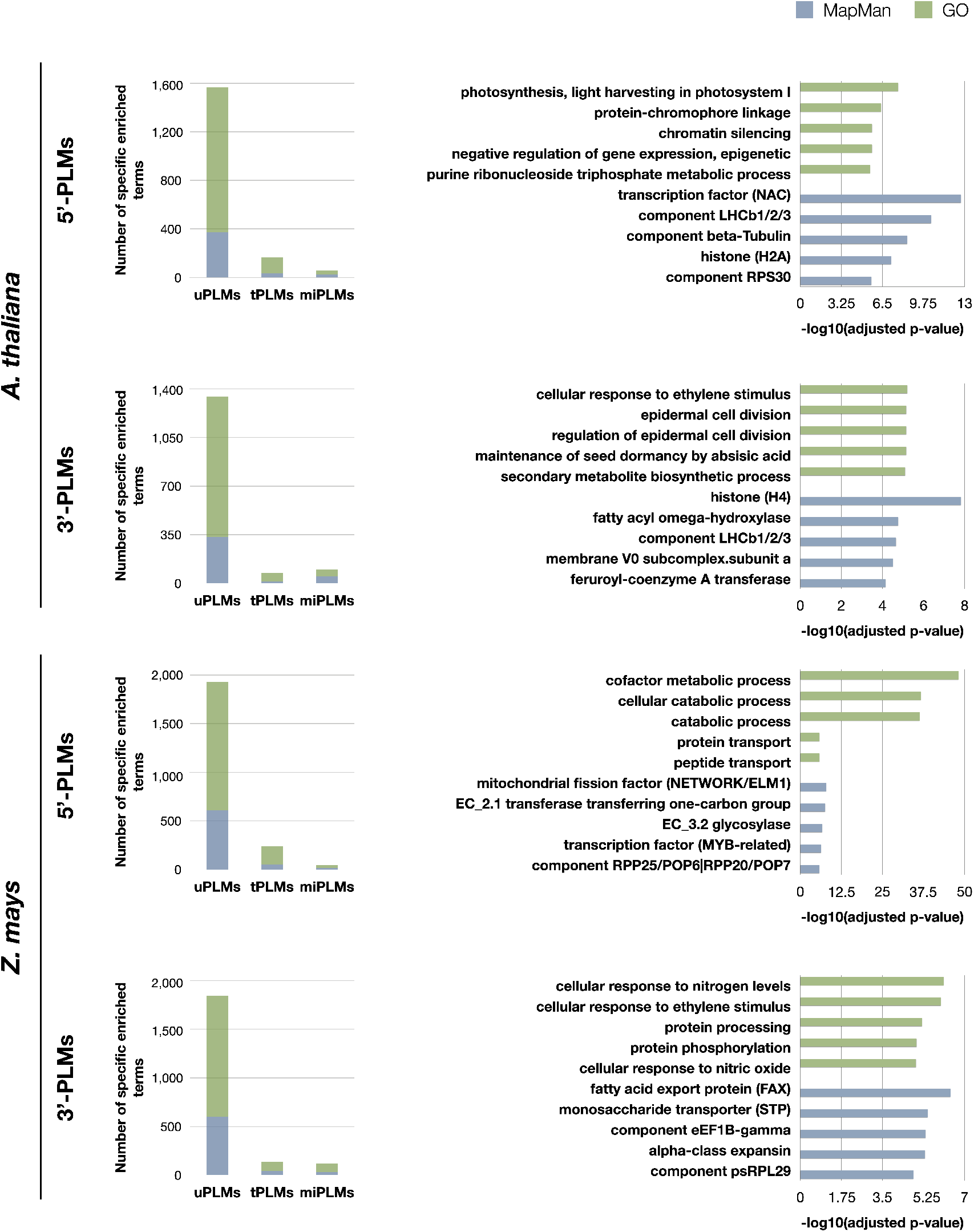
GO and MapMan terms enriched specifically for each type of PLMs-containing gene sets. Values for GO and Mapman terms are shown in green and blue, respectively. On the left: histograms of the number of GO and MapMan terms enriched specifically for each type of PLMs-containing gene sets in the two species studied. On the right: the 5 most enriched terms specifically for uPLMs. The bar values of the histograms indicate the -log(adjusted p-values) for each term. MapMan terms have been truncated at the maximum precision level. The corresponding integer MapMan terms are as follows: transcription factor (NAC): RNA biosynthesis.transcriptional regulation.transcription factor (NAC); component LHCb1/2/3: Photosynthesis.photophosphorylation.photosystem II.LHC-II complex.component LHCb1/2/3; component beta-Tubulin: Cytoskeleton organisation.microtubular network.alpha-beta-Tubulin heterodimer.component beta-Tubulin; histone (H2A): Chromatin organization.histones.histone (H2A); component RPS30: Protein biosynthesis.ribosome biogenesis.small ribosomal subunit (SSU).SSU proteome.component RPS30; histone (H4): Chromatin organisation.histones.histone (H4); fatty acyl omega-hydroxylase: Cell wall organisation.cutin and suberin.cuticular lipid formation.fatty acyl omega-hydroxylase; membrane V0 subcomplex.subunit a: Solute transport.primary active transport.V-type ATPase complex.membrane V0 subcomplex.subunit a; feruroyl-coenzyme A transferase: Cell wall organisation.cutin and suberin.alkyl-hydrocinnamate biosynthesis.feruroyl-coenzyme A transferase; mitochondrial fission factor (NETWORK/ELM1): Cell cycle organisation.organelle division.mitochondrion and peroxisome division.mitochondrial fission factor (NETWORK/ELM1); EC_2.1 transferase transferring one-carbon group: Enzyme classification.EC_2 transferases.EC_2.1 transferase transferring one-carbon group; EC_3.2 glycosylase: Enzyme classification.EC_3 hydrolases.EC_3.2 glycosylase; transcription factor (MYB-related): RNA biosynthesis.transcriptional regulation.MYB transcription factor superfamily.transcription factor (MYB-related); component RPP25/POP6|RPP20/POP7: RNA processing.ribonuclease activities.RNA-dependent RNase P complex.component RPP25/POP6|RPP20/POP7; fatty acid export protein (FAX): Lipid metabolism.lipid trafficking.fatty acid export protein (FAX); monosaccharide transporter (STP): Solute transport.carrier-mediated transport.MFS superfamily.SP family.monosaccharide transporter (STP); component eEF1B-gamma: Protein biosynthesis.translation elongation.eEF1 aminoacyl-tRNA binding factor activity.eEF1B eEF1A-GDP-recycling complex.component eEF1B-gamma; alpha-class expansin: Cell wall organisation.cell wall proteins.expansin activities.alpha-class expansin; component psRPL29 Protein biosynthesis.organelle machinery.plastidial ribosome.large ribosomal subunit proteome.component psRPL29.

Comparing 5’- and 3’-uPLMs-containing gene sets revealed specific terms associated with each of the two sets (Supplementary Fig. 2 and Table 7). Notably, we observed that “cellular response to ethylene stimulus” was one of the five most enriched GO terms in the 3’-uPLMs-containing gene set of both *A. thaliana* and *Z. mays*. Some of the genes considered are characterized by uPLMs signals in the −450 to −200 bases relative to the TTS. These signals are further supported by the fact that the uPLM sequences are very conserved between the two species (CGTCG and its reverse-complementary CGACG for *A. thaliana*; ACGCCCAC / GGGCGTCC and its reverse-complementary GGACGCCC for *Z. mays*). Terms related to “Cell wall organisation” were also present in the five most enriched MapMan terms for the *A. thaliana*-uPLMs-contaigning gene set in both regions, although the protein classes identified were different. For example, we found that part of the genes encoding alpha-expansin are characterized by 5’-uPLMs signals localized after the TSS, while part of the genes encoding acyl omega-hydroxylases are characterized by 3’-uPLMs signals localized in group 1 and 2.

As expected, each species had also specific enriched terms (Supplementary Fig. 2 and Table 7). For the 5’-uPLMs-gene set, these terms were mainly related to “cell killing” in *A. thaliana,* while they were associated with “transposition” and antibiotic metabolic/catabolic processes in *Z. mays* (Supplementary Table 8g-8h). For the 3’-uPLMs-gene set, specific enriched terms were once again related to “cell killing” in *A. thaliana,* while they were mainly related to “translation” in *Z. mays*. Taken together, these findings reveal that uPLMs provide functional predictions distinct from those inferred by tPLMs and miPLMs.

### Biological processes associated to uncharacterized genes through integration of PLM information

Taking into account the contribution of PLMs, we thought to use them to infer gene regulatory networks and go further and deeper into the characterization of some gene families (Fig. 1). We focused on a poorly characterized, *Z. mays*-specific gene family (referred to as HOM04M002476 by PLAZA) defined only by the GO term “transposition” (Supplementary Table 8g). This gene family consists of 65 genes, 64 of which were considered in the detection of 5’-PLMs (Supplementary Table 11a). Using the 5’-tPLMs detected for all these 64 genes, we investigated the TF-target gene relationships (Fig. 7a). A total of 545 tPLMs were associated with 416 TFs belonging to 37 distinct TF families. Among them, AP2/ERF domain, Myb-related and WRKY were the three most abundant TF families (Supplementary Table 11).

**Fig. 7.**
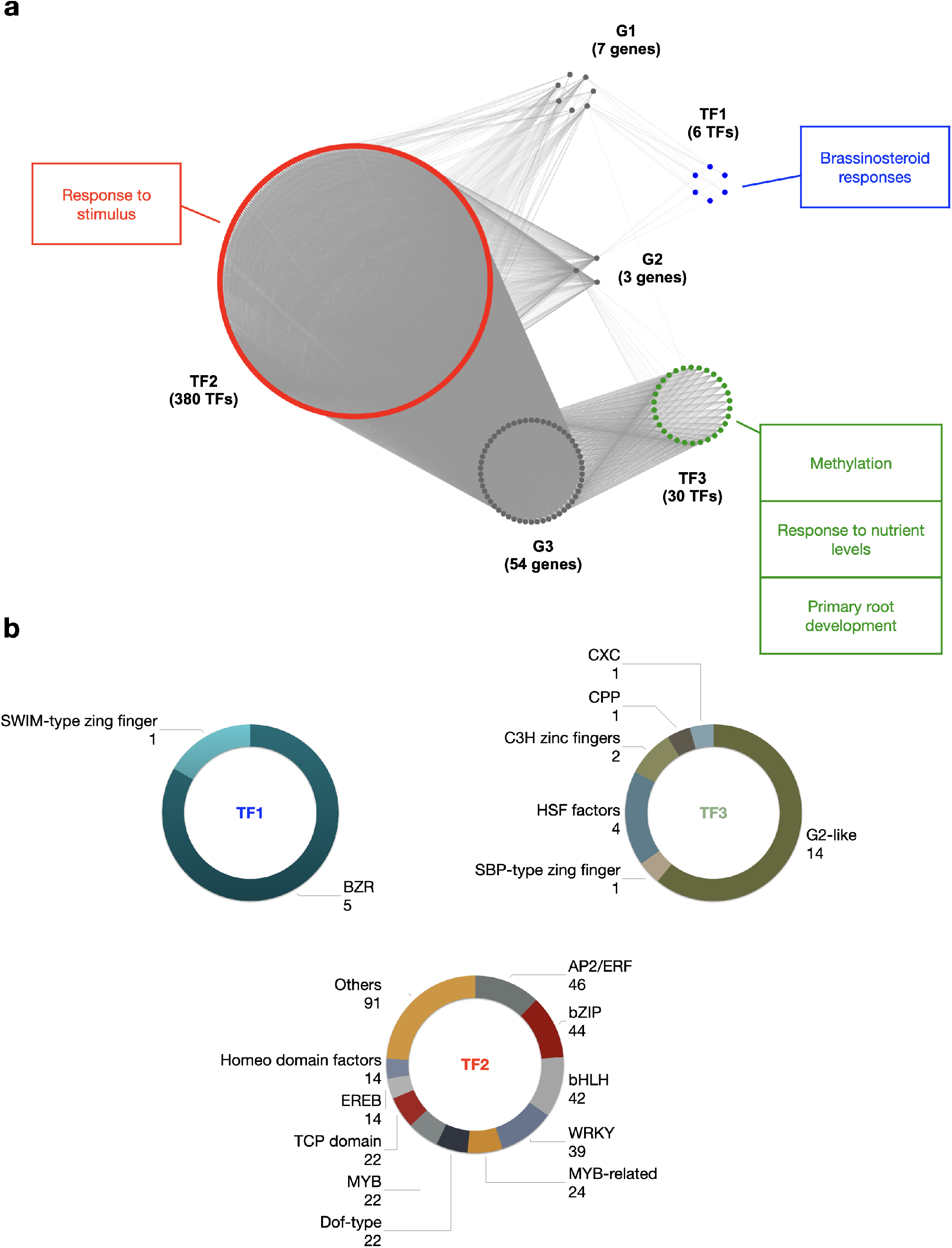
Gene regulatory network of the HOM04M002476 gene family of *Z. mays*. (**a**) Topological structure of the gene regulatory network of the HOM04M002476 gene family. G1, G2 and G3 are the three HOM04M002476 gene modules (grey) identified by the LBM procedure. TF1 (red), TF2 (green) and TF3 (blue) are the three TF modules identified by the LBM procedure potentially involved in the regulation of HOM04M002476 genes. GO enrichments of TF modules are indicated with corresponding colors. (**b**) TF families associated to the three TF modules.

Clustering based on latent block model (LBM) revealed three modules of target genes referred to as G1, G2 and G3 with 7, 3 and 54 members, respectively (Fig. 7a and Supplementary Table 11). It also revealed three modules of TFs referred to as TF1, TF2 and TF3 with 6, 380 and 30 TFs belonging to 2, 29 and 6 TF families, respectively (Fig. 7a–7b and Supplementary Table 11). We found that genes belonging to module G1 were regulated by TFs from modules TF1 (2/2 families), TF2 (16/29 families) and TF3 (1/6 family). It is worth noting that the SWIM-type zinc finger TF family of module TF1 was specific to genes from G1 module. Genes belonging to module G2 were also regulated by TFs belonging to all three modules, including the BZR TF family of module TF1, all TF families of module TF2, and the HSF factors and G2-like TF families of module TF3. In contrast, genes belonging to module G3 were only regulated by TFs from modules TF2 and TF3. Furthermore, the CXC, CPP and SBP-type zinc finger TF families of module TF3 covered specifically genes from G3 module.

To elucidate the potential involvement of this gene regulatory network in biological processes, we conducted GO and MapMan enrichment analysis of the TF modules. As expected, terms enriched in a common way in all three modules were related to the regulation of transcription (Supplementary Table 12). Additionally, in module TF1, the most specifically enriched terms were related to brassinosteroid responses (Fig. 7a and Supplementary Table 12a-12b). In module TF2, except for terms related to transcriptional regulation, the most enriched terms was related to “response to stimulus” (Fig. 7a; Supplementary Table 12c-12d). Finally, in module TF3 the most enriched terms were mainly related to methylation, response to nutrient levels and primary root development (Fig. 7a and Supplementary Table 12e-12f). Our computational approach therefore paves the way for pinpointing the function of this gene family in these distinct processes in future follow-up studies.

## Discussion

Understanding gene transcriptional regulation requires understanding where regulatory factors bind genomics DNA. Although several efforts have recently been undertaken to characterize TFBSs, the identification of high resolution *cis*-regulatory elements at the genome-wide scale remains an arduous challenge. Hence, we attempted to reveal the whole PLM landscape by using large-scale and *de novo* PLM detection to systematically profile proximal *cis*-regulatory elements. The three PLM group structure revealed in *A. thaliana* and *Z. mays* with distantly related genomes echoes and enriches established knowledge of the 5’-gene-proximal region (Fig. 8). Omitting potential annotation errors, we found that the localization of these motifs, including the core promoter, was less constrained in *Z. mays* compared to *A. thaliana*, as recently reported for TATA-boxes at varying distances from TSS in *Z. mays* (42). Similarly, we observed that the dispersion of tPLMs remained more important in *Z. mays* than in *A. thaliana*, indicating that putative TFBSs have also a less constrained preferential localization in *Z. mays* than in *A. thaliana.* Overall, these data suggest that the 5’-proximal genomic context may be less constrained in *Z. mays* than in *A. thaliana*. This might be related to the richness of the *Z. mays* genome in transposable elements compared to that of *A. thaliana* (43).

**Fig. 8.**
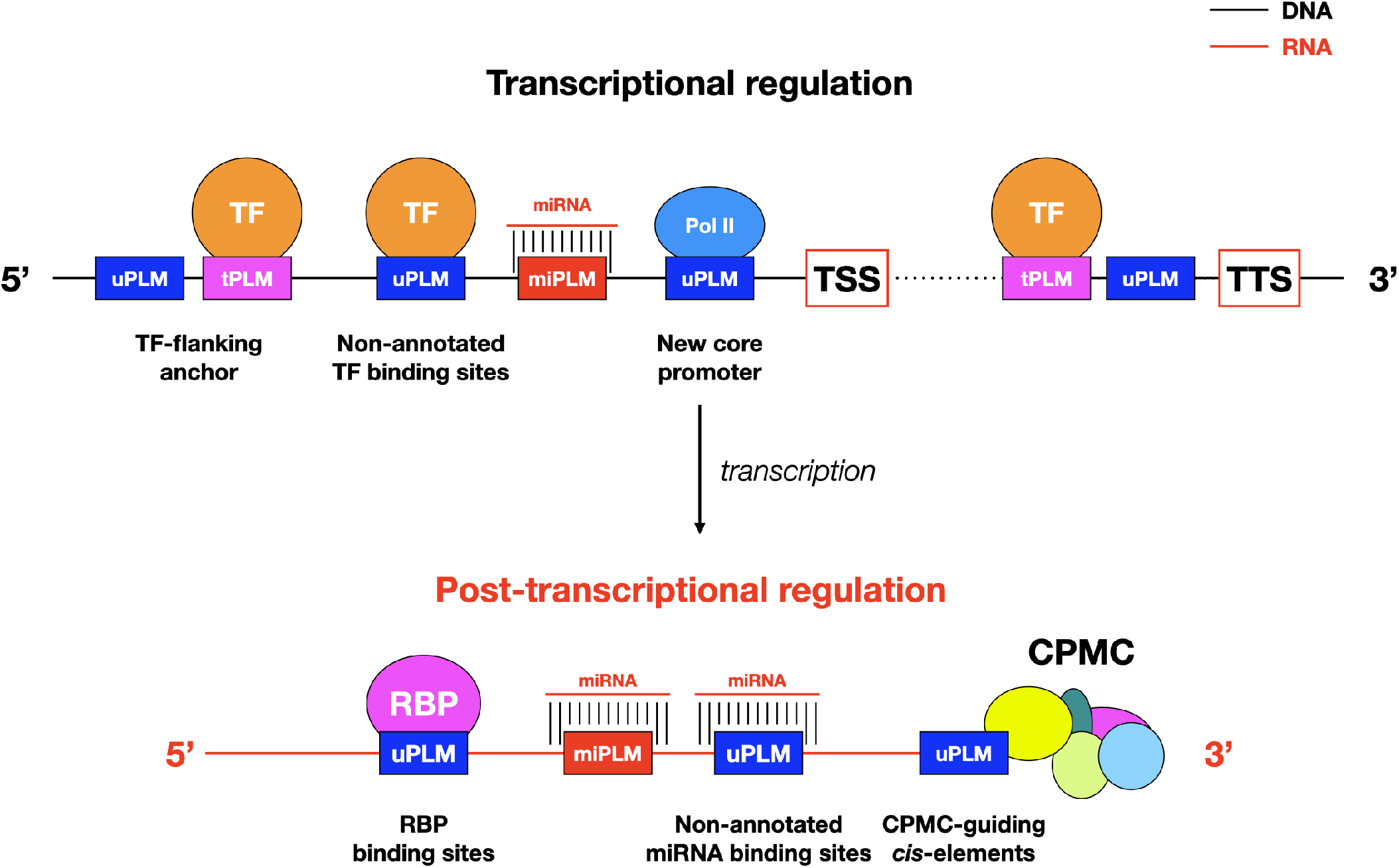
Models of PLM-mediated transcriptional and post-translational controls in plants. The models integrate only the results presented here. Possible interactions between the illustrated components and the diverse array of other intermediates such as enhancers and long-distance chromatin interaction sites, remain to be investigated. CPMC: cleavage and polyadenylation molecular complex; Pol II: RNA polymerase II; RBP: RNA-binding proteins.

Our finding of conserved PLMs beween *A. thaliana* and *Z. mays* suggests that the closer we get to the genes, the more the context, including *cis*-regulatory elements (here given by tPLMs), are conserved between species. Notably, we showed that this context seems to be more conserved in the 3’-proximal region than in the 5’-proximal region of genes, emphasizing the importance of the 3’-gene-proximal region in genomic structure. Despite its key role in gene expression, the 3’-gene-proximal region remains poorly studied in plants (25, 30, 44). Because the density maxima observed for tPLMs and uPLMs was reached in the *cis*-elements that guide the CPMC and overlapped with groups 2 and 3, it is quite possible that the 3’-PLMs detected in these portions of DNA sequence constitute a catalog of NUE, FUE and CE (Fig. 8). In support of this hypothesis, we observed that the AATAAA motif (and its complementary reverse), which is the key site involved in polyadenylation and is extremely conserved in mammals and somewhat less in plants, was located between 10 and 20 bases upstream of the TTS. Furthermore, the nucleotide percentages of 3’-PLMs detected in these regulatory regions are consistent with known proportions of nucleotides in these *cis*-elements. Together, these data support the idea that 3’-PLMs may constitute an accurate catalog of CPMC-guiding *cis*-elements. In this respect, the presence of 3’-tPLMs in this catalog (around 11%) and more generally in the whole region, opens interesting mechanistic perspectives on the role of TFs. First, they may act as activators or repressors of the transcriptional machinery (Fig. 8). Second, by binding to tPLMs located in the FUE/NUE and CE regions, they could impact pre-mRNAs length and thus mRNA stability by influencing the choice of an alternative polyadenylation site at the end of transcripts (25, 30, 44).

The retained proportion observed between uPLMs (up to 25 %) and tPLMs (up to 30 %), also suggests that uPLMs could constitute a context that needs to be conserved. In this regard, the uPLM genomic significance is already supported by experimental MOA-seq data that validate some of the uPLMs identified. This raises the question of what mechanisms underlie the presence of uPLMs. First, as mentionned earlier, some uPLMs may be players in the core promoter or polyadenylation process (Fig. 8). Second, some uPLMs may be non-annotated binding sites. Indeed, we showed that the 104 maize TF ChIP-seq data (32) contributed 16% and 33% more tPLMs for *A. thaliana* and *Z. mays,* respectively, compared to the JASPAR Plant 2020 database that was updated prior to the release of these ChIP-seq data. Similar to the post-transcriptional regulation by miRNAs, RNA-binding proteins (45, 46) are major players that can potentially bind PLMs at the transcriptional level. Consequently, there is no doubt that future resources will supplement the assignment of uPLMs. Finally, uPLMs may be motifs that are not directly bound by TFs but that play a crucial role in the correct binding of these regulators to neighboring TFBSs (Fig. 8) (47–49). This concept of “flanking sequence context” appears extremely relevant because of the nature of PLMs, which are constrained motifs at a distance from genes. This idea also raises many questions about the existence and role of tPLMs/uPLMs associations, with tPLMs bound by TFs and uPLMs serving as essential context sequences for the formation of the DNA-TF complex. Additional analyses and integration with other *in vivo* information will be key to advance functional tests needed to ascertain the relative importance of tPLMs and uPLMs as *cis*-regulatory elements controlling gene expression. Meanwhile, our results have broader implications for future characterization of unannotated genes in plants.

## Methods

### Genomic datasets

TAIR10 (50) and B73v4.39 (51) genomes and their annotations were considered to extract the 5’- and 3’-gene-proximal sequences of *A. thaliana* and *Z. mays* genes, respectively.

### Preparation of the gene-proximal sequence files

For the 5’-gene-proximal region, annotation of the TSS was ensured by filtering genes without a 5’-UTR region (in GFF3/GTF file). Genes on reverse strand were reverse-complemented to analyze all sequences in the same orientation. Extracted sequences corresponded to the intervals [-1000;+500] and [-1500;+500] bp relative to the TSS for *A. thaliana* and *Z. mays,* respectively. In total, 19,736 and 25,848 genes were analyzed for *A. thaliana* and *Z. mays*, respectively. For the 3’-gene-proximal region, similarly to what has been done for the 5’-gene-proximal region sequences, annotation of the TTS was ensured by filtering genes without a 3’-UTR region annotated. To standardize the PLM detection step, genes on forward strand were reverse-complemented. Extracted sequences were [-500;+1000] and [-500;+1500] bp with respect to the TTS for *A. thaliana* and *Z. mays*, respectively. Taking in consideration only annotated 3’-UTR, 20,573 and 25,199 genes were processed for *A. thaliana* and *Z. mays*, respectively.

### Preparation of the motif file

Every non-polymorphic DNA 4-mers to 8-mers was generated representing 87,296 motifs. Among these motifs, 256, 1,024, 4,096, 16,384, and 65,536 had a length of 4, 5, 6, 7 and 8 bp, respectively.

### Large-scale and *de novo* PLM detection

To reveal PLMs in each gene-proximal region of each species, we calculated the number of the motif occurrences at each position to get a motif distribution as described previously (20). First, the gene-proximal sequence file and the motif file were given as inputs. Second, a linear regression was computed on a neutral region defined as the first (last) 500 bp for the 5’ (3’)-gene-proximal region where no accumulation of PLMs is expected. Third, predicted values were calculated in the studied region defined as the gene-proximal region with a confidence interval of 99%. If the observed occurence distribution exceeded the confidence interval of the prediction in the studied region, the motif was considered as a PLM. The latter is characterized by (i) its preferential position, defined as the position where the distribution exceeds the confidence interval and the maximum high is reached, (ii) a functional window, defined as the part of the studied region where the motif distribution exceeds the confidence interval, and (iii) a score defined as the difference between the high of the peak and the upper bound of the confidence interval at the preferential position. To check PLM redundancy, we calculated the Jaccard index of each pair of PLMs for each PLM containing-gene set with an inclusion link. This index was obtained by dividing the intersection of the two lists by their union. A Jaccard index of 1 indicates a perfect match between the gene lists. This index was also calculated on the functional window of each pair of PLMs to quantify their overlap.

### TFBS and microRNA resources and assignment

TF-BSs (676 total) were extracted from JASPAR Plant 2020 (31) and ChIP-seq of 104 maize leaf TFs (32). MicroRNAs (miRNAs) were obtained from psRNATarget (37) and only those from *A. thaliana* (427 total) and *Z. mays* (321 total) were kept.

We first assigned TFBSs to PLMs for both species in each region separately using the TOMTOM web tool (52). Euclidean distance was next used as a comparison function with a q-value threshold at 0.05 and the complete scoring option deselected. PLMs were also compared to the top 1% of k-mers of the 104 maize TFs (32) by considering only exact matches. Because miRNAs regulate genes by sequence complementarity (35) and are molecules from 19 to 22 nt, we only considered our 8 bp PLMs for this comparison (the 20 bp size by which miRNAs regulate gene expression was not tested due to computational time constraints). If a PLM was exactly found in a miRNA, it was assigned as a miPLM.

### Functional annotation

Functional annotation of genes from both species was based on MapMan X4 (53) and Gene Ontology (GO) from PLAZA 4.5 (54). Functional enrichment analysis of genes containing an identified PLM (Supplementary Datasets) was performed by comparing the relative occurrence of each term to its relative occurrence in a reference list for each region and species using a hypergeometric test with the R function phyper. These reference lists consisted of all genes considered for PLM detection in each gene-proximal region in both species as described in the ‘Preparation of the gene-proximal sequence files’ of the Methods section. P-values were adjusted by the Benjamini-Hochberg (BH) procedure to control the False Discovery Rate (FDR). An enriched term had its adjusted P-value lower than 0.05.

### Comparative analysis of PLMs and MOA-seq motifs

*Z. mays* 5’-PLMs were compared to published MOA-seq motifs (16) using TOMTOM with the same parameters as described for TFBSs assignment and according to the following two criteria: (1) both sequence types had to be identical (TOMTOM q-value < 0.05) and (2) the position of the MOA-seq motif had to be within the functional window of the PLM extended by 50 bases upstream and downstream.

### Inference and topology analysis of the HOM04M002476 gene family regulatory network

A matrix of dimension 37×64 was generated with TF families in rows and HOM04M002476 genes in columns. Links between TF families and target genes were established when tPLMs, and thus associated TFs, were identified for a given target gene. These links were indicated by ones in the matrix. A zero indicated the absence of a tPLM associated with the TF family in the target gene. Gene and TF modules were obtained using LBM with the R package blockmodels (55).

Functional enrichment of each TF module was performed by comparing the relative occurrence of each term to its relative occurrence in the list of genes encoding TFs in *A. thaliana* (2,208 genes) and *Z. mays* (2,164 genes) using a hypergeometric test with the R phyper function. *P*-values were adjusted by the BH procedure to control the FDR. An enriched term had its adjusted *P*-value lower than 0.05.

## Acknowledgements

We thank all members of the ‘Genomic Networks’ (GNet) and ‘Biomass Quality and Interactions with Drought’ (QUALIBIOSEC) teams past and present. We also thank Hank W. Bass for sharing some works prior to publication and Hervé Vaucheret for very helpful discussions. This work was supported by the Plant2Pro® Carnot Institute in the frame of the PLMViewer program. Plant2Pro® is supported by ANR (agreement #19 CARN 0024 01). The IPS2 and IJPB laboratories benefit from the support of Saclay Plant Sciences-SPS (ANR-17-EUR-0007). J.R. is supported by a PhD fellowship of the Doctoral School ‘Structure and Dynamics of Living Systems’ from the University Paris-Saclay.

## Data availability

All the data supporting this work are available in Supplementary data.

## Code availability

The codes used to generate the data are available at https://forgemia.inra.fr/GNet/plmdetect/plmdetect_tool.

## Author contributions

V.B, M-L.M. and S.C. conceived the project. M-L.M. and S.C. designed and supervised the study. J.R. generated, analyzed and interpreted the data. C.G. formatted the functional annotation files. J.R., M-L.M. and S.C. wrote the manuscript. All the authors read the paper and agree with the final version.

