## Supplementary Figures for "A comprehensive map of preferentially located motifs reveals distinct proximal *cis*-regulatory elements in plants"

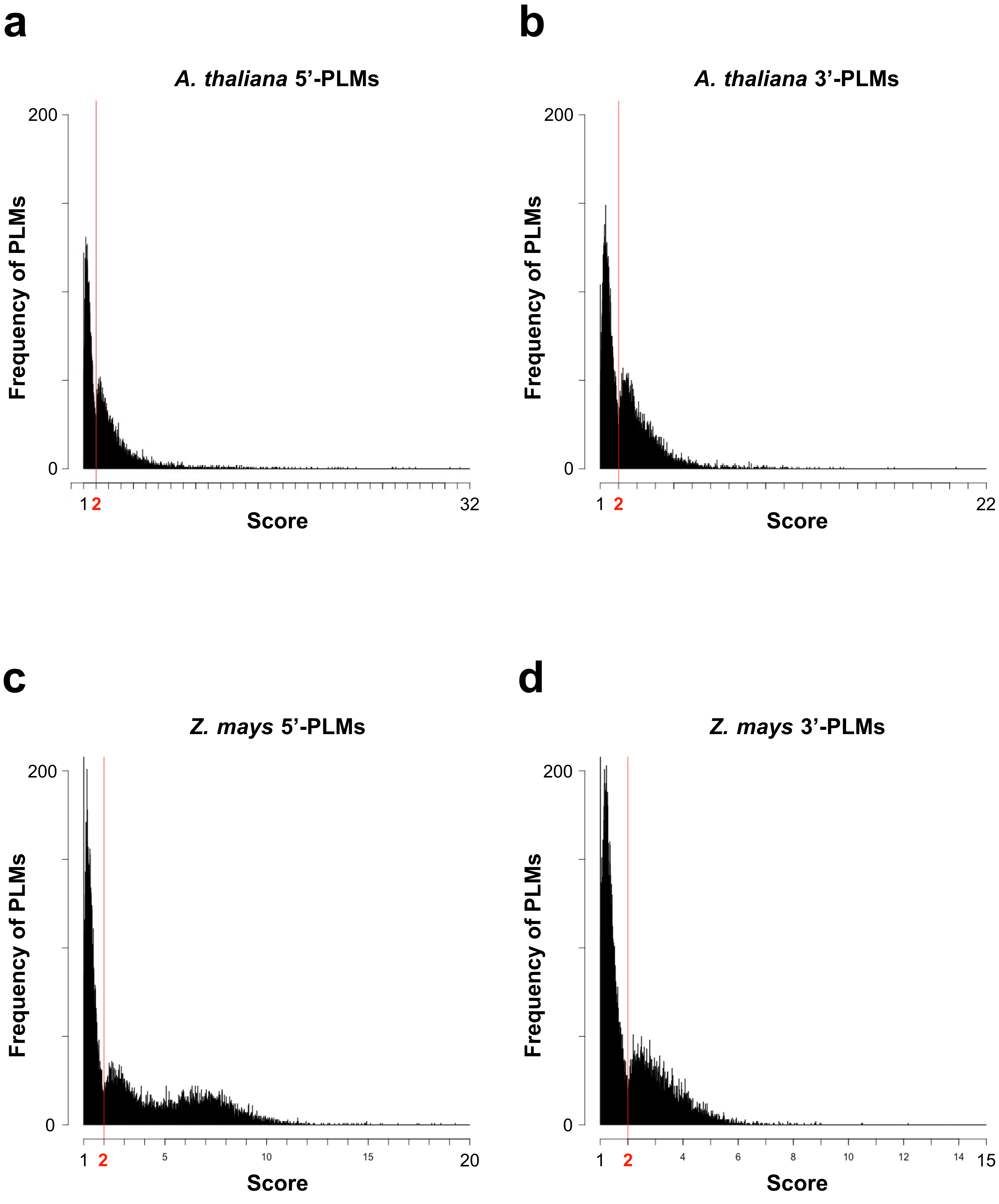


**Supplementary Figure 1.** Histograms of PLM frequency relative to their score. The red line indicates the separation in the distribution at the score of 2. **(a-c)** Number of 5'-PLMs relative to the score in *A. thaliana* **(a)** and *Z. mays* **(c)**. **(b-d)** Number of 3'-PLMs relative to the score in *A. thaliana* **(b)** and *Z. mays* **(d)**.

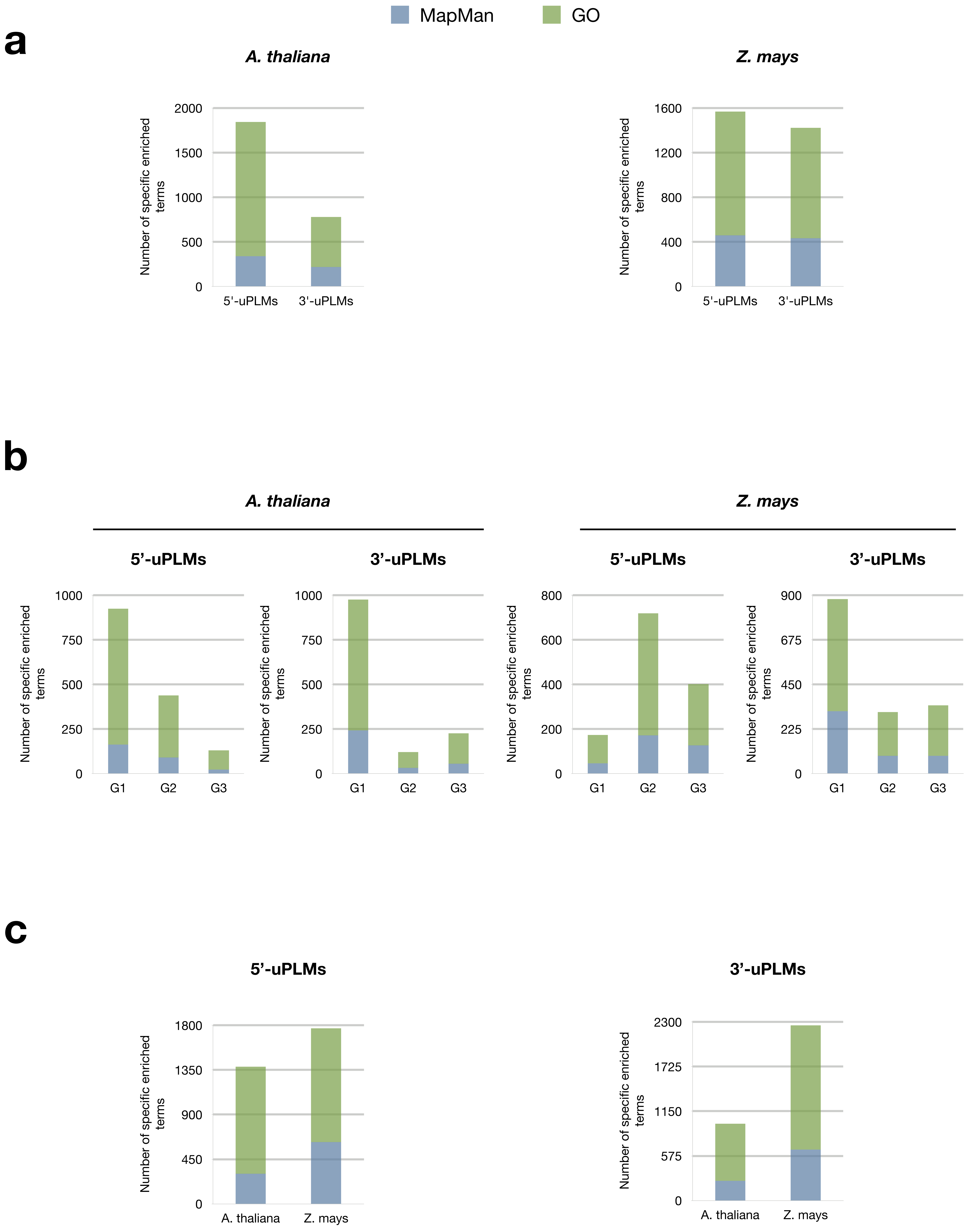


**Supplementary Figure 2.** Dissecting specific enriched GO and MapMan terms in uPLMs-containing gene sets in *A. thaliana* and *Z. mays*. **(a)** Number of specific enriched terms in each uPLMs-containing gene set compared to the other set in each species. **(b)** Number of specific enriched terms for each group of uPLMs-containing gene for both regions and species. **(c)** Number of specific enriched terms in the uPLMs-containing gene set of *A. thaliana* for each proximal-region compared to that of *Z. mays* for the same proximal region.
