## Supplementary material for "A comprehensive map of preferentially located motifs reveals distinct proximal *cis*-regulatory elements in plants": ATAAGCGG_containing_gene_set.txt

#Occurences of ATAAGCGG between -159 and -151 (positions relative to the transcription start site)
#15 sequences with ATAAGCGG in this interval
#0.06% of the sequences in promoter set with ATAAGCGG (set with 25199 sequences)
Zm00001d047605::9:136921777-136923778(-)
Zm00001d024910::10:94434703-94436704(+)
Zm00001d047811::9:142368703-142370704(+)
Zm00001d047809::9:142343263-142345264(+)
Zm00001d018952::7:10372255-10374256(-)
Zm00001d047810::9:142360087-142362088(-)
Zm00001d034222::1:287044875-287046876(+)
Zm00001d044865::9:5840797-5842798(-)
Zm00001d053118::4:214483747-214485748(+)
Zm00001d051510::4:160973658-160975659(+)
Zm00001d010923::8:133673022-133675023(-)
Zm00001d052830::4:202078494-202080495(-)
Zm00001d038035::6:145770269-145772270(+)
Zm00001d005024::2:154433882-154435883(+)
Zm00001d008765::8:19527779-19529780(-)
