## Supplementary material for "A comprehensive map of preferentially located motifs reveals distinct proximal *cis*-regulatory elements in plants": ATAGGCGA_containing_gene_set.txt

#Occurences of ATAGGCGA between -1000 and -981 (positions relative to the transcription start site)
#13 sequences with ATAGGCGA in this interval
#0.05% of the sequences in promoter set with ATAGGCGA (set with 25199 sequences)
Zm00001d053524::4:233017449-233019450(+)
Zm00001d048017::9:148140138-148142139(-)
Zm00001d052992::4:207904167-207906168(+)
Zm00001d035515::6:30768714-30770715(-)
Zm00001d010196::8:103510146-103512147(+)
Zm00001d028751::1:45511787-45513788(-)
Zm00001d039007::6:168670052-168672053(+)
Zm00001d009562::8:70205311-70207312(-)
Zm00001d037956::6:143536005-143538006(+)
Zm00001d049880::4:50008394-50010395(+)
Zm00001d031057::1:175081054-175083055(-)
Zm00001d030697::1:154087090-154089091(+)
Zm00001d025700::10:126771226-126773227(+)
