## Supplementary material for "A comprehensive map of preferentially located motifs reveals distinct proximal *cis*-regulatory elements in plants": CATACGTT_containing_gene_set.txt

#Occurences of CATACGTT between -319 and -288 (positions relative to the transcription start site)
#24 sequences with CATACGTT in this interval
#0.10% of the sequences in promoter set with CATACGTT (set with 25199 sequences)
Zm00001d024854::10:90670913-90672914(-)
Zm00001d019914::7:75385692-75387693(-)
Zm00001d019108::7:17234844-17236845(-)
Zm00001d032727::1:236334095-236336096(+)
Zm00001d050640::4:108479145-108481146(-)
Zm00001d038955::6:167630686-167632687(+)
Zm00001d043442::3:200057507-200059508(+)
Zm00001d045621::9:29035885-29037886(+)
Zm00001d021371::7:150145139-150147140(+)
Zm00001d003758::2:57751492-57753493(-)
Zm00001d047010::9:115073020-115075021(-)
Zm00001d004965::2:150854066-150856067(-)
Zm00001d027861::1:15585485-15587486(-)
Zm00001d019957::7:80230684-80232685(+)
Zm00001d042308::3:160128634-160130635(+)
Zm00001d020004::7:85029696-85031697(+)
Zm00001d049824::4:46624381-46626382(+)
Zm00001d038648::6:161706362-161708363(-)
Zm00001d004333::2:104142859-104144860(+)
Zm00001d023621::10:12505142-12507143(-)
Zm00001d040971::3:85996865-85998866(+)
Zm00001d004936::2:149584438-149586439(+)
Zm00001d041208::3:105546943-105548944(-)
Zm00001d051332::4:154910690-154912691(+)
