## Supplementary material for "A comprehensive map of preferentially located motifs reveals distinct proximal *cis*-regulatory elements in plants": CCAACGTA_containing_gene_set.txt

#Occurences of CCAACGTA between -331 and -282 (positions relative to the transcription start site)
#24 sequences with CCAACGTA in this interval
#0.10% of the sequences in promoter set with CCAACGTA (set with 25199 sequences)
Zm00001d036827::6:102931889-102933890(+)
Zm00001d047958::9:146173388-146175389(+)
Zm00001d007430::2:231537183-231539184(-)
Zm00001d032993::1:246129730-246131731(+)
Zm00001d019914::7:75385692-75387693(-)
Zm00001d047010::9:115073020-115075021(-)
Zm00001d009997::8:93872991-93874992(+)
Zm00001d003758::2:57751492-57753493(-)
Zm00001d045384::9:20250431-20252432(-)
Zm00001d049824::4:46624381-46626382(+)
Zm00001d040971::3:85996865-85998866(+)
Zm00001d053687::4:239035150-239037151(+)
Zm00001d023402::10:4739146-4741147(+)
Zm00001d029898::1:93138276-93140277(-)
Zm00001d013140::5:5354345-5356346(+)
Zm00001d004965::2:150854066-150856067(-)
Zm00001d027861::1:15585485-15587486(-)
Zm00001d050640::4:108479145-108481146(-)
Zm00001d012888::5:1614748-1616749(-)
Zm00001d011697::8:159026631-159028632(+)
Zm00001d051332::4:154910690-154912691(+)
Zm00001d041208::3:105546943-105548944(-)
Zm00001d004333::2:104142859-104144860(+)
Zm00001d020891::7:135607560-135609561(+)
