## Supplementary material for "A comprehensive map of preferentially located motifs reveals distinct proximal *cis*-regulatory elements in plants": CGGCACAA_containing_gene_set.txt

#Occurences of CGGCACAA between 347 and 360 (positions relative to the transcription start site)
#24 sequences with CGGCACAA in this interval
#0.10% of the sequences in promoter set with CGGCACAA (set with 25199 sequences)
Zm00001d016154::5:147355331-147357332(+)
Zm00001d009753::8:79786965-79788966(+)
Zm00001d022608::7:181004811-181006812(+)
Zm00001d033859::1:276071834-276073835(+)
Zm00001d046941::9:111968501-111970502(-)
Zm00001d004619::2:122403987-122405988(+)
Zm00001d039356::3:2464720-2466721(+)
Zm00001d024822::10:89616473-89618474(+)
Zm00001d016048::5:141503341-141505342(+)
Zm00001d027747::1:12533690-12535691(+)
Zm00001d046296::9:78818408-78820409(+)
Zm00001d032393::1:225188462-225190463(-)
Zm00001d031576::1:195076207-195078208(+)
Zm00001d010426::8:114525754-114527755(+)
Zm00001d010871::8:131613912-131615913(-)
Zm00001d004047::2:78778958-78780959(+)
Zm00001d010293::8:107993794-107995795(-)
Zm00001d016486::5:164511395-164513396(-)
Zm00001d009297::8:52933440-52935441(-)
Zm00001d006513::2:210304554-210306555(+)
Zm00001d006615::2:212638440-212640441(-)
Zm00001d010148::8:101476363-101478364(+)
Zm00001d051764::4:168965668-168967669(+)
Zm00001d023220::10:698908-700909(+)
