## Supplementary material for "A comprehensive map of preferentially located motifs reveals distinct proximal *cis*-regulatory elements in plants": CGTGTTAG_containing_gene_set.txt

#Occurences of CGTGTTAG between -994 and -946 (positions relative to the transcription start site)
#20 sequences with CGTGTTAG in this interval
#0.08% of the sequences in promoter set with CGTGTTAG (set with 25199 sequences)
Zm00001d013447::5:11829362-11831363(-)
Zm00001d018030::5:212392781-212394782(-)
Zm00001d052450::4:190300219-190302220(+)
Zm00001d027842::1:14880957-14882958(+)
Zm00001d012015::8:166512540-166514541(+)
Zm00001d028457::1:35509530-35511531(-)
Zm00001d013758::5:19600076-19602077(-)
Zm00001d020965::7:138122417-138124418(-)
Zm00001d038470::6:157953486-157955487(+)
Zm00001d037540::6:128728306-128730307(-)
Zm00001d016928::5:180246096-180248097(+)
Zm00001d046468::9:91363829-91365830(+)
Zm00001d028509::1:37480611-37482612(-)
Zm00001d030051::1:102221404-102223405(-)
Zm00001d052903::4:204209240-204211241(-)
Zm00001d043139::3:190001016-190003017(+)
Zm00001d052905::4:204262166-204264167(-)
Zm00001d033879::1:277216123-277218124(+)
Zm00001d016821::5:177259272-177261273(-)
Zm00001d006677::2:214085348-214087349(+)
