## Supplementary material for "A comprehensive map of preferentially located motifs reveals distinct proximal *cis*-regulatory elements in plants": GCAGTAAG_containing_gene_set.txt

#Occurences of GCAGTAAG between 183 and 193 (positions relative to the transcription start site)
#21 sequences with GCAGTAAG in this interval
#0.08% of the sequences in promoter set with GCAGTAAG (set with 25199 sequences)
Zm00001d039934::3:19864379-19866380(+)
Zm00001d029354::1:67217126-67219127(-)
Zm00001d031925::1:206933997-206935998(-)
Zm00001d052536::4:192537679-192539680(-)
Zm00001d047054::9:116803863-116805864(+)
Zm00001d047400::9:128764840-128766841(-)
Zm00001d018763::7:4524133-4526134(+)
Zm00001d022609::7:181033305-181035306(-)
Zm00001d012866::5:1362449-1364450(-)
Zm00001d028632::1:41459256-41461257(+)
Zm00001d003705::2:55528558-55530559(-)
Zm00001d021075::7:142290627-142292628(-)
Zm00001d016980::5:181879364-181881365(-)
Zm00001d016206::5:150428182-150430183(+)
Zm00001d018765::7:4584872-4586873(+)
Zm00001d028298::1:29616883-29618884(-)
Zm00001d029694::1:82891721-82893722(+)
Zm00001d044044::3:217676618-217678619(-)
Zm00001d015191::5:78810026-78812027(+)
Zm00001d031920::1:206691935-206693936(+)
Zm00001d011155::8:140993541-140995542(+)
