## Supplementary material for "A comprehensive map of preferentially located motifs reveals distinct proximal *cis*-regulatory elements in plants": GCTGCCCG_containing_gene_set.txt

#Occurences of GCTGCCCG between 394 and 403 (positions relative to the transcription start site)
#19 sequences with GCTGCCCG in this interval
#0.08% of the sequences in promoter set with GCTGCCCG (set with 25199 sequences)
Zm00001d004910::2:148017214-148019215(-)
Zm00001d032942::1:244209891-244211892(-)
Zm00001d052595::4:194256227-194258228(+)
Zm00001d017334::5:193208445-193210446(+)
Zm00001d021070::7:142138300-142140301(-)
Zm00001d023396::10:4672874-4674875(+)
Zm00001d033130::1:251634226-251636227(+)
Zm00001d041491::3:123758700-123760701(+)
Zm00001d043179::3:190704269-190706270(-)
Zm00001d042146::3:153570290-153572291(-)
Zm00001d035772::6:46914143-46916144(+)
Zm00001d017335::5:193224198-193226199(+)
Zm00001d039155::6:171180492-171182493(-)
Zm00001d048066::9:149281934-149283935(+)
Zm00001d002139::2:6551371-6553372(-)
Zm00001d013053::5:3892563-3894564(-)
Zm00001d009660::8:74949844-74951845(+)
Zm00001d016682::5:172406584-172408585(+)
Zm00001d019669::7:49323361-49325362(-)
