## Supplementary material for "A comprehensive map of preferentially located motifs reveals distinct proximal *cis*-regulatory elements in plants": GTCACTGT_containing_gene_set.txt

#Occurences of GTCACTGT between -131 and -124 (positions relative to the transcription start site)
#20 sequences with GTCACTGT in this interval
#0.08% of the sequences in promoter set with GTCACTGT (set with 25199 sequences)
Zm00001d043819::3:210961866-210963867(+)
Zm00001d003896::2:65737733-65739734(+)
Zm00001d029648::1:81311546-81313547(-)
Zm00001d042515::3:170298863-170300864(+)
Zm00001d016711::5:173430742-173432743(-)
Zm00001d015737::5:114785384-114787385(+)
Zm00001d039503::3:6082514-6084515(-)
Zm00001d012950::5:2321180-2323181(-)
Zm00001d044993::9:9497746-9499747(-)
Zm00001d033808::1:274300169-274302170(+)
Zm00001d027709::1:11677945-11679946(-)
Zm00001d016968::5:181545374-181547375(-)
Zm00001d049376::4:28202325-28204326(+)
Zm00001d054098::4:246454035-246456036(+)
Zm00001d015455::5:91168860-91170861(-)
Zm00001d044559::3:231693124-231695125(+)
Zm00001d015672::5:105096523-105098524(+)
Zm00001d019619::7:46403812-46405813(-)
Zm00001d047929::9:145593270-145595271(-)
Zm00001d028520::1:38039114-38041115(-)
