## Supplementary material for "A comprehensive map of preferentially located motifs reveals distinct proximal *cis*-regulatory elements in plants": GTGGTCCG_containing_gene_set.txt

#Occurences of GTGGTCCG between 435 and 446 (positions relative to the transcription start site)
#18 sequences with GTGGTCCG in this interval
#0.07% of the sequences in promoter set with GTGGTCCG (set with 25199 sequences)
Zm00001d005761::2:187001549-187003550(-)
Zm00001d031920::1:206691935-206693936(+)
Zm00001d032949::1:244465005-244467006(+)
Zm00001d028632::1:41459256-41461257(+)
Zm00001d018763::7:4524133-4526134(+)
Zm00001d044044::3:217676618-217678619(-)
Zm00001d032954::1:244533638-244535639(+)
Zm00001d003705::2:55528558-55530559(-)
Zm00001d015191::5:78810026-78812027(+)
Zm00001d006719::2:215151606-215153607(-)
Zm00001d018761::7:4473391-4475392(+)
Zm00001d018765::7:4584872-4586873(+)
Zm00001d043148::3:190151073-190153074(+)
Zm00001d018408::5:220204595-220206596(-)
Zm00001d023600::10:11479067-11481068(+)
Zm00001d052536::4:192537679-192539680(-)
Zm00001d002119::2:6131807-6133808(-)
Zm00001d021249::7:146835547-146837548(-)
