## Supplementary material for "A comprehensive map of preferentially located motifs reveals distinct proximal *cis*-regulatory elements in plants": TGTCGCAG_containing_gene_set.txt

#Occurences of TGTCGCAG between 397 and 408 (positions relative to the transcription start site)
#20 sequences with TGTCGCAG in this interval
#0.08% of the sequences in promoter set with TGTCGCAG (set with 25199 sequences)
Zm00001d026083::10:138057458-138059459(-)
Zm00001d025874::10:132246654-132248655(-)
Zm00001d050133::4:68207702-68209703(+)
Zm00001d035386::6:23945743-23947744(+)
Zm00001d032688::1:234472697-234474698(+)
Zm00001d003935::2:70121007-70123008(+)
Zm00001d019432::7:33468517-33470518(+)
Zm00001d038306::6:154121526-154123527(-)
Zm00001d005530::2:177343813-177345814(+)
Zm00001d023934::10:29996129-29998130(+)
Zm00001d046493::9:92348033-92350034(+)
Zm00001d048908::4:7745546-7747547(+)
Zm00001d049865::4:49255762-49257763(-)
Zm00001d041926::3:143906674-143908675(-)
Zm00001d025047::10:101524452-101526453(+)
Zm00001d044290::3:224370067-224372068(-)
Zm00001d027520::1:7358098-7360099(-)
Zm00001d052570::4:193375059-193377060(-)
Zm00001d031099::1:177233414-177235415(-)
Zm00001d002632::2:17556997-17558998(+)
