## Supplementary material for "A comprehensive map of preferentially located motifs reveals distinct proximal *cis*-regulatory elements in plants": TTAATACG_containing_gene_set.txt

#Occurences of TTAATACG between 0 and 23 (positions relative to the transcription start site)
#21 sequences with TTAATACG in this interval
#0.08% of the sequences in promoter set with TTAATACG (set with 25199 sequences)
Zm00001d011704::8:159245369-159247370(-)
Zm00001d020288::7:104311270-104313271(+)
Zm00001d023899::10:27171147-27173148(-)
Zm00001d038206::6:151542733-151544734(+)
Zm00001d020916::7:136511655-136513656(+)
Zm00001d016214::5:150603375-150605376(-)
Zm00001d046710::9:103065705-103067706(-)
Zm00001d002974::2:28499282-28501283(-)
Zm00001d042313::3:160479170-160481171(+)
Zm00001d044152::3:220527271-220529272(-)
Zm00001d053340::4:226746641-226748642(-)
Zm00001d019163::7:19753823-19755824(-)
Zm00001d042973::3:185494565-185496566(-)
Zm00001d028285::1:29169823-29171824(+)
Zm00001d043437::3:199879556-199881557(-)
Zm00001d002388::2:11652379-11654380(+)
Zm00001d023537::10:9346464-9348465(-)
Zm00001d053277::4:223882107-223884108(+)
Zm00001d011173::8:141782767-141784768(-)
Zm00001d021502::7:154176211-154178212(-)
Zm00001d047681::9:139140882-139142883(+)
